# Mitochondrial small RNA alterations associated with increased lysosome activity in an Alzheimer’s Disease Mouse Model uncovered by PANDORA-seq

**DOI:** 10.1101/2024.10.18.619155

**Authors:** Xudong Zhang, Junchao Shi, Pratish Thakore, Albert L. Gonzales, Scott Earley, Qi Chen, Tong Zhou, Yumei Feng Earley

## Abstract

Emerging small noncoding RNAs (sncRNAs), including tRNA-derived small RNAs (tsRNAs) and rRNA-derived small RNAs (rsRNAs), are critical in various biological processes, such as neurological diseases. Traditional sncRNA-sequencing (seq) protocols often miss these sncRNAs due to their modifications, such as internal and terminal modifications, that can interfere with sequencing. We recently developed panoramic RNA display by overcoming RNA modification aborted sequencing (PANDORA-seq), a method enabling comprehensive detection of modified sncRNAs by overcoming the RNA modifications. Using PANDORA-seq, we revealed a novel sncRNA profile enriched by tsRNAs/rsRNAs in the mouse prefrontal cortex and found a significant downregulation of mitochondrial tsRNAs and rsRNAs in an Alzheimer’s disease (AD) mouse model compared to wild-type controls, while this pattern is not present in the genomic tsRNAs and rsRNAs. Moreover, our integrated analysis of gene expression and sncRNA profiles reveals that those downregulated mitochondrial sncRNAs negatively correlate with enhanced lysosomal activity, suggesting a crucial interplay between mitochondrial RNA dynamics and lysosomal function in AD. Given the versatile tsRNA/tsRNA molecular actions in cellular regulation, our data provide insights for future mechanistic study of AD with potential therapeutic strategies.

**Graphical Abstract:** 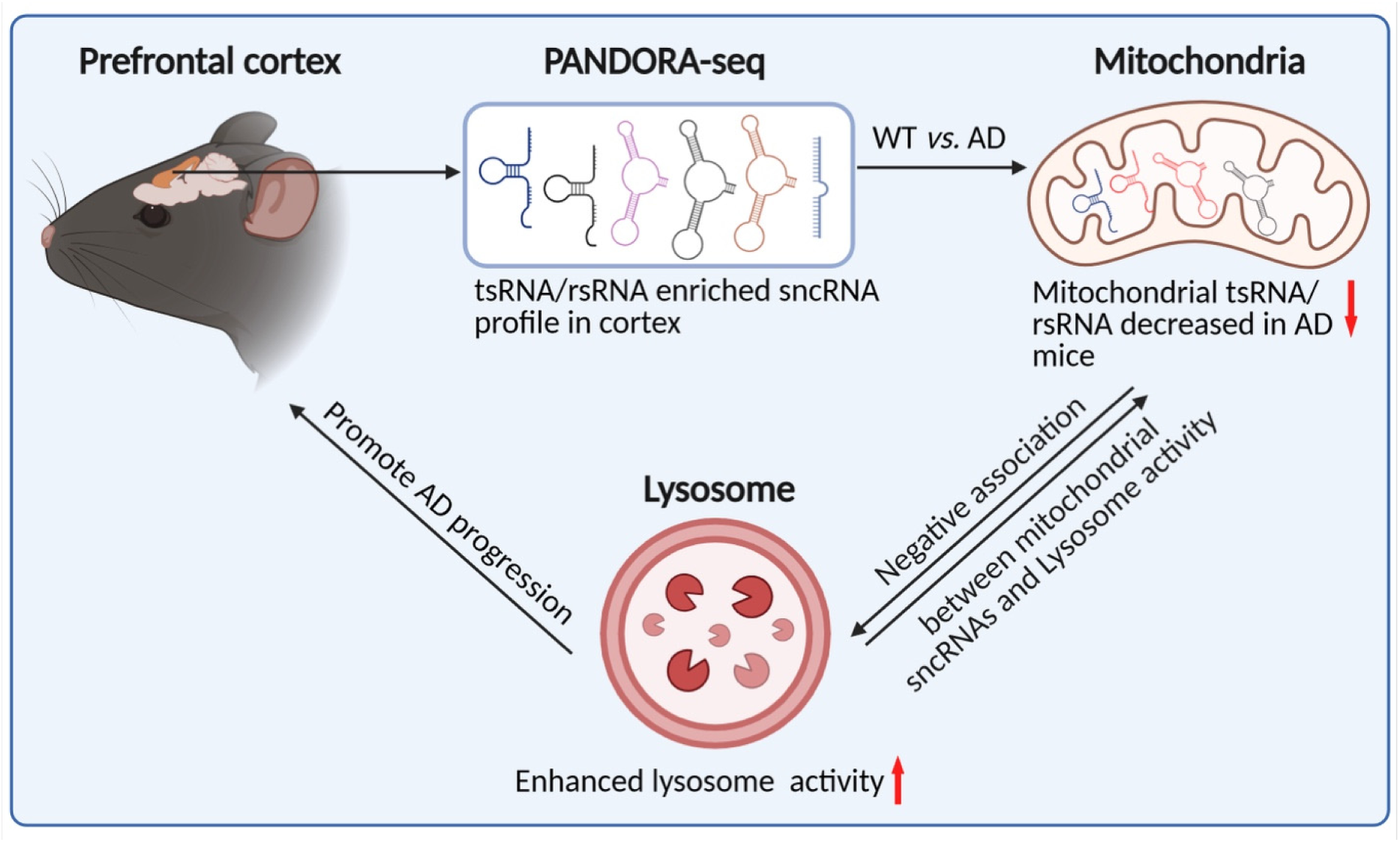

## 1. Introduction

Alzheimer’s disease (AD) is the predominant cause of all dementias, accounting for 60%-80% of cases. Most AD cases are sporadic with an unknown etiology, whereas very few AD patients have familial AD (FAD), which is inherited in an autosomal dominant manner^1^. AD is neuropathologically characterized by the accumulation of toxic β-amyloid protein (Aβ) proteins and hyperphosphorylated tau proteins, leading to cognitive impairments. Both accumulations are closely linked with the progression of AD. A recent study in a human cohort demonstrated that FAD gene mutations influence the accumulation patterns of Aβ peptides, contributing to the heterogeneity of Aβ plaque characteristics among patients at similar stages of AD progression^2^. The 5XFAD mice model, developed by introducing three *App* gene mutations (KM670/671NL, I76V, V717I) alongside two *Psen1* mutations (M146L, L286V) into transgenic mice^3^ has been widely adopted in AD research. Although hyperphosphorylated tau and neurofibrillary tangles may be absent, this model has been proven to successfully recapitulate key pathological features of AD, including the formation of Aβ plaques and loss of synapses and neurons. In addition, 5XFAD mice show Aβ-dependent and age-dependent progressive deficits in spatial and learning memory, alongside motor impairments at an elder age^4^. Despite advances in understanding the genetic and environmental factors driving AD, many aspects of AD remain unclear. One emerging area of interest is the role of small noncoding RNAs (sncRNAs) in AD progression, particularly their influence on gene regulation and neurodegeneration.

Small non-coding RNAs (sncRNAs), particularly the emerging tRNA-derived small RNAs (tsRNAs) and rRNA-derived small RNAs (rsRNAs) have been shown to play important roles in diverse biological processes, including stem cell maintenance^5, 6^, tumorigenesis^7, 8^, epigenetic inheritance^9–12^, viral infections^13, 14^, and the pathogenesis in neurological diseases^15, 16^. However, the traditional sncRNA-seq methods have limitations in comprehensively detecting highly modified sncRNAs such as tsRNAs and rsRNAs, as their internal and terminal modifications can interfere with cDNA library construction procedures, leading to substantial biases where these modified sncRNAs cannot be efficiently included in the sequencing results. To address these issues, we have recently developed PANDORA-seq, a novel sequencing method that overcomes the limitations in traditional sncRNA sequencing by addressing specific RNA modifications and sncRNA termini, enabling comprehensive analysis of previously undetectable sncRNAs, particularly tsRNAs and rsRNA carrying these RNA modifications and terimini, leading to a more comprehensive of sncRNA landscape in the studied tissue and cell types^17^.

In this report, we profiled sncRNA landscape in prefrontal cortex samples of Alzheimer’s disease mice (AD) mice (5XFAD mice) and control mice by using PANDORA-seq, revealing an unprecedented sncRNA landscape compared to the use of traditional sncRNA-seq, and found that the downregulated mitochondrial sncRNAs are negatively correlated with enhanced lysosomal activity in the AD model, suggesting that these mitochondrial tsRNAs are potential novel factors involved in the pathogenesis and/or progression of Alzheimer’s disease.

## 2. Methods and materials

### 2.1 Animals and behavior test

#### Mice

Hemizygous 5XFAD mice and WT littermates of both sexes on a congenic C57BL6/J background (MMRCC stock #34848) were used^18^. All animal care procedures and experimental protocols involving animals complied with the NIH Guide for the Care and Use of Laboratory Animals and were approved by the Institutional Animal Care and Use Committees at the University of Nevada, Reno. Mice were maintained in individually ventilated cages (<5 mice/cage) with ad libitum access to food and water in a room with controlled 12-h light and dark cycles.

#### Y-Maze Behavioral Assay

The spontaneous alternation behavioral test was employed to assess short-term spatial working memory using the Y-maze^19, 20^. Mice were placed in one of the three maze arms (the start arm) and allowed to explore all three arms for 10 minutes. The sessions were recorded and analyzed using EthoVision XT software (version 16.0.1536, Noldus Information Technology, Leesburg, VA, USA). Spontaneous alternation was assessed by tracking the sequence of entries into each arm during the 10-minute session. The alternation index (%) was calculated as (number of spontaneous alternations / maximum possible alternations) × 100. Spontaneous alternation was defined as consecutive entries into each maze arm without repetition, and the maximum alternation was the total possible alternations (total number of arm entries – 2).

#### Novel Object Recognition Test (NORT)

Memory recall was assessed using the Novel Object Recognition Test (NORT), as described previously^21^. The test was conducted over three days, consisting of habituation, training, and testing phases. On Day 1, mice were habituated by being placed in an empty arena and allowed to explore for 10 minutes. On Day 2, during the training phase, two identical objects were placed in the arena. The mice were then placed in the center of the arena and allowed to explore both objects for 10 minutes. On Day 3, during the testing phase, one of the objects from the training phase (the familiar object) was replaced with a novel object. Mice were placed into the center of the arena and allowed to explore for 10 minutes. Sessions were recorded and analyzed using EthoVision XT software (Noldus Information Technology). The recognition index (%) was calculated as (total exploration time for the novel object during testing / total exploration time for the familiar object during testing) × 100.

### 2.2 Isolation of specified-size small RNAs

Total RNA was isolated from 100 mg of mice prefrontal tissues using the TRIzol reagent (Invitrogen™), as described by the manufacturer. Then, the RNA sample, mixed with an equal volume of 2× RNA loading dye (New England Biolabs), was incubated at 75 °C for 5 min. The mixture was loaded into 15% (wt/vol) Urea-polyacrylamide gel, followed by electrophoresis in 1× TBE running buffer at 200 V for 40 minutes. After staining with SYBR Gold solution (Invitrogen; S11494), gels that contained small RNAs of 15–50 nucleotides were excised based on small RNA ladders (Abnova^TM^ R0007). Small RNAs were recovered from the gel as previously described. In short, the gel was immersed in 0.3 M sodium acetate solution containing 20 Unit RNase inhibitor (New England Biolabs, M0314L) overnight at 4 °C. The sample was centrifuged for 10 min at 12,000g (4 °C). The supernatant was mixed with 3 volumes of ethanol, 1/10 volume of 3 M sodium acetate, and 1ul linear acrylamide. Then, the sample was incubated at −20 °C for 2 h and centrifuged for 25 min at 12,000g. After removing the supernatant, the precipitation was resuspended in nuclease-free water, quantified, and stored at −80 °C until analysis.

### 2.3 Small RNA sequencing by PANDORA-seq and traditional-seq

PANDORA-Seq protocol has been recently described in our report^17^. In short, the sncRNA isolated from the prefrontal cortex for PANDORA-seq will go through Alkane monooxygenase (AlkB) and T4 polynucleotide kinase (T4 PNK) treatment steps before library construction by NEBNext® Small RNA Library Prep Kit. The sncRNA for traditional-seq directly went to the library construction workflow without enzyme treatment. The workflow of small RNA library construction was conducted as the manufacturer described.

For AlkB enzyme treatment, the sncRNA was incubated in a 50 μl reaction mixture containing 50 mM HEPES, 75 μM ferrous ammonium sulfate, 1 mM α-ketoglutaric acid, 2 mM sodium ascorbate, 50 mg/L BSA, 200 ng ALKB enzyme, 20 Unit RNase inhibitor at 37°C for 30 min. Then, the mixture was added into 500 μl TRIzol reagent for RNA isolation. For T4 PNK enzyme treatment, sncRNA was incubated in 50 μl reaction mixture containing 5 μl 10× PNK buffer, 1 mM ATP, 10 U T4 PNK (New England Biolabs) at 37°C for 20 min. Then, the mixture was added to 500 μl TRIzol reagent for RNA isolation. All cDNA were sequenced on Illumina^TM^ Novaseq 6000 platform.

### 2.4 Small RNA expression analysis

The raw data (i.e., the .fastq files) from both PANDORA-seq and traditional sncRNA-seq were analyzed and annotated by the software *SPORTS1.1*^22^. We then employed the *edgeR* algorithm to perform the pairwise comparison in sncRNA expression between groups. The *TMM* algorithm was applied to perform read count normalization, and a likelihood ratio test was employed to identify differentially expressed sncRNAs. The sncRNA species with a false discovery rate (*FDR*) <0.05 and fold change (*FC*) >2 was defined as differentially expressed.

### 2.5 Transcriptome and pathway analysis

The creation of cDNA libraries and sequencing were performed by the Illumina standard workflow. mRNA expression level was quantified using the *Salmon* tool with default setting. We then employed the *edgeR* algorithm to compare the transcriptomic pattern between groups. We applied the TMM algorithm for read count normalization and likelihood ratio test for differentially expressed gene (DEG) identification. The genes with *FDR* <0.05 and *FC* >1.5 were deemed differentially expressed. We further performed pathway analysis upon the DEGs based on the KEGG pathways, using the David tool^23, 24^. In addition, for each KEGG pathway, we computed a gene-set score, using the Functional Analysis of Individual Microarray Expression (*FAIME*) algorithm^25^. A higher *FAIME* score suggests an overall upregulation of a given pathway^25^. The gene expression heatmap was generated using the “heatmap.2” function within the “gplots” R package. The hierarchical clustering was performed using the “complete” method with “euclidean” distance. The co-expression between “Lysosome” pathway and sncRNA families was performed using *Spearman*’s rank correlation test.

### 2.6 Statistical analysis

All the statistical analyses were conducted using the R programming platform and Prism8 software (GraphPad, La Jolla, CA, USA). Data are presented as means ± SEM except for the high-throughput sequencing data. Student t-test or Ordinary One-way ANOVA with post hoc Bonferroni correction was used as appropriate for comparisons among groups. A P-value < 0.05 was considered statistically significant. In the case of multiple testing, P-value adjustment was performed using the “p.adjust” function with the “BH” method in R programming.

## 3. Results

### 3.1 Alzheimer’s disease mice model shows cognitive impairment

To evaluate and confirm the cognitive impairments of Alzheimer’s disease (AD) mouse model (5xFAD), we conducted the Y-maze assay, a tool for assessing short-term memory^26^. The spontaneous alternation percentage was measured at both the two-month and six-month stages. Typically, mice without cognitive deficits prefer exploring less recently visited arms of the Y-maze, resulting in a higher spontaneous alternation percentage (Fig. 1A). At two-months-old, there was no significant difference in alternation index between WT and AD groups (Fig. 1B). In contrast, at the six-month stage, AD mice showed incorrect spontaneous alternation and a lower alternation index compared to the wildtype (WT) mice indicating reduced spatial working memory (Fig. 1B). Additionally, there were no notable differences in maximum alternation (Fig. 1C) between the WT and AD mice at either the two-month or six-month stage showing the total number of alternations possible was the same between the two groups.

**Fig. 1.**
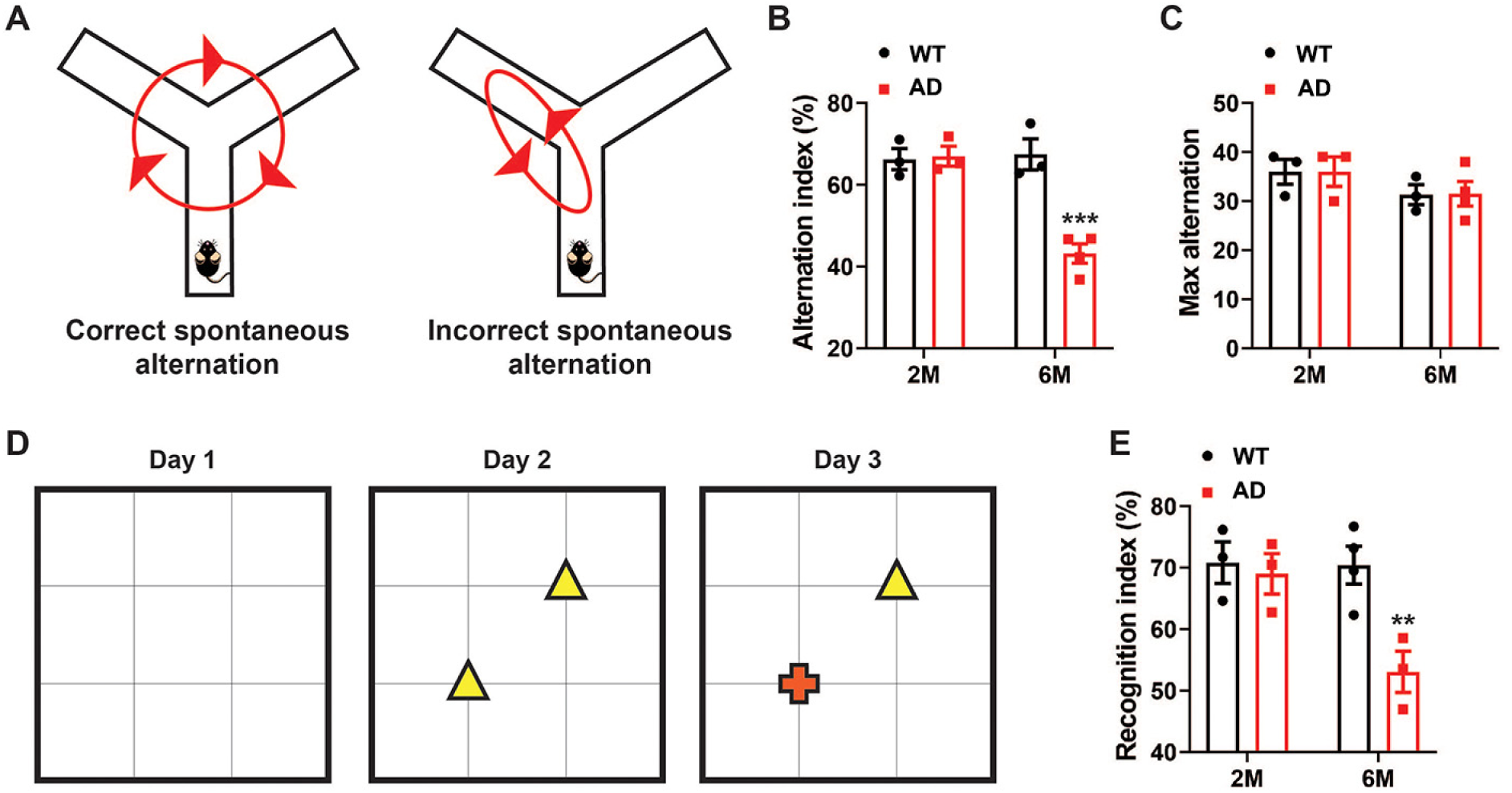
Age-dependent cognitive impairment in transgenic AD mice. (A) Schematic of the Y-maze Spontaneous alternation behavior assay showing an example of a correct (left) and incorrect (right) spontaneous alternation. (B) The alternation index in the Y-maze test in two- and six-month-old WT and AD mice (***p<0.001, n=3-4 in each group). (C) Maximum alternation by WT and AD mice in Y-maze test at two-month and six-month age, respectively. (D) Schematic of Novel Object Recognition test (NORT) protocol consisting of a habituation phase (Day 1), training phase (Day 2), and testing phase (Day 3). A novel object is introduced during the testing phase, and the time spent exploring the novel object is assessed. (E) The recognition index during the testing phase in two- and six-month-old WT and AD mice (**p<0.01, n=3-4 in each group). Data was analyzed by two-way ANOVA with Sidak post-test for multiple comparisons.

Next, we utilized the Novel Object Recognition Test (NORT), which is based on innate preference of mice for novel objects^26^, to evaluate recognition memory between WT and AD mice (Fig. 1D). AD mice at the six-month stage spent less time exploring a novel red crossed-shaped object compared to WT mice (Fig. 1D), with a recognition index of approximately 53%, which is significantly lower than the 70% observed in WT mice (Fig.1E). Again, we didn’t observe significant difference between AD mice and WT mice at the two-month stage in the NORT task assay. These results suggest that the 5xFAD mice mimic aspects of cognitive impairment observed in Alzheimer’s disease.

### 3.2 Pandora-seq shows an unprecedented sncRNAs landscape enriched by tsRNAs and rsRNAs in the prefrontal cortex of mice

To determine PANDORA-seq’s efficacy in uncovering modified sncRNA populations within neural tissues, we processed RNAs from the cortex of WT and AD mice using both traditional sncRNA seq and PANDORA-seq followed by the bioinformatic analysis with SPORTS 1.1 pipeline, an optimized tool for small RNA sequencing data analysis^22^. Traditional sequencing predominantly revealed a miRNA-enriched sncRNA landscape in the cortex in which miRNAs account for 56% of the total small RNA reads identified (Fig. 2A). This finding is consistent with the expectation that miRNAs are readily captured by traditional sequencing protocol due to their hypomodification nature. In contrast, PANDORA-seq uncovered a sncRNA landscape enriched with tsRNAs and rsRNAs, with tsRNAs (26.7%) and rsRNAs (35.6%) together accounting for 62.3% of the total sncRNAs reads (Fig. 2B). Moreover, PADORA-seq demonstrated a notable proficiency in capturing piRNA population within the classical 26-31nt length range, while the traditional-seq majorly detected piRNA population in a shorter 21-24nt length range (Fig. 2). These results highlight the advantage of PADORA-seq in providing a more comprehensive and accurate sncRNA landscape in mice prefrontal cortex. We next further analyzed the sncRNAs landscape of AD and WT mice cortex via PANDORA-seq data, the result consistently highlighted the ability of PANDORA-seq to delineate a tsRNA/rsRNA enriched sncRNA landscape. Notably, we observed a decrease of 44-nt-length rsRNAs in the AD mice group, in both the males and females, compared to the WT mice (Supplementary Fig.1).

**Fig. 2.**
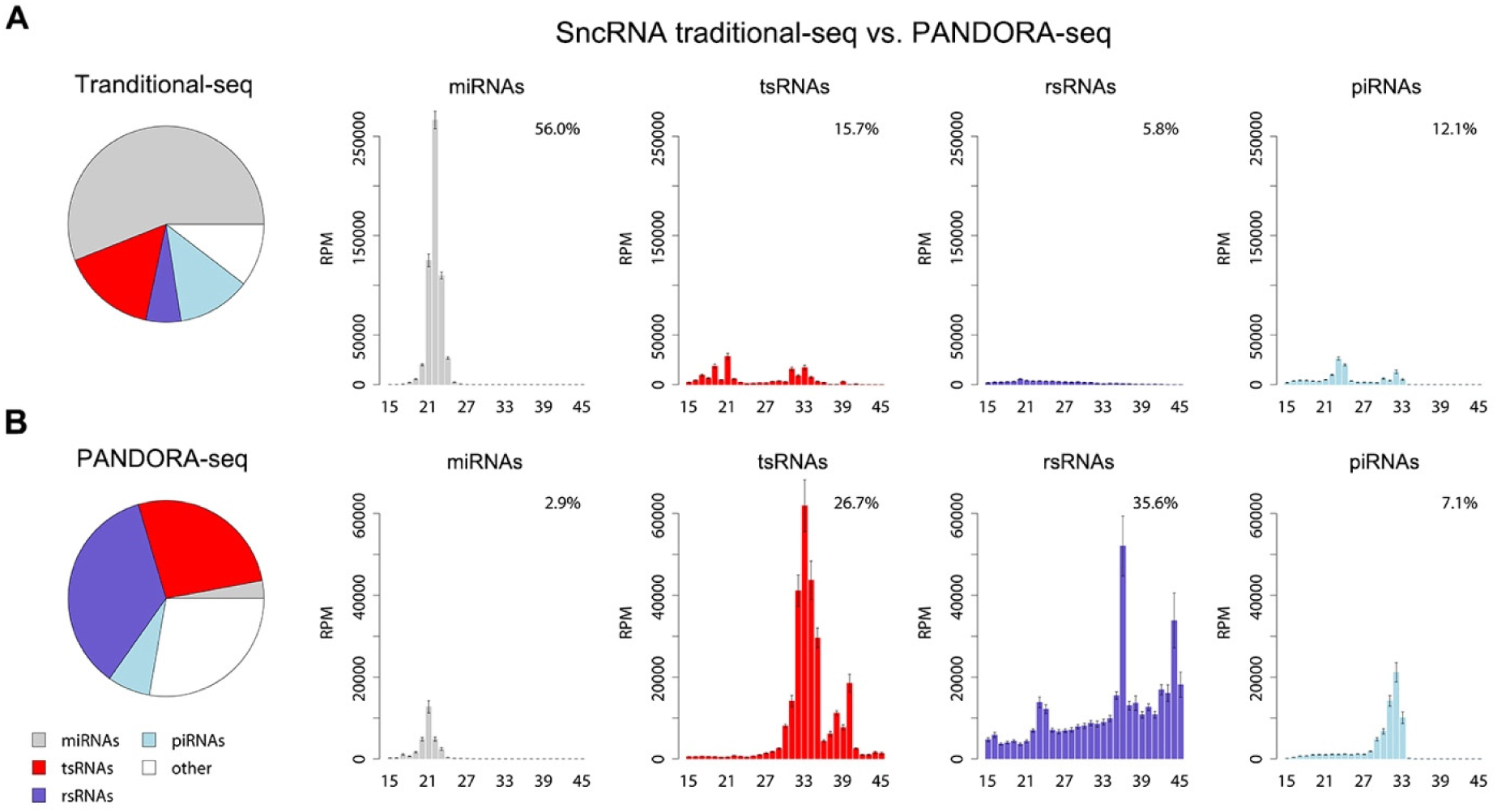
PANDORA-seq revealed a tsRNA/rsRNA-dominant sncRNA landscape in the prefrontal cortex. Representative result of sncRNAs species detected by traditional-seq and PANDORA-seq respectively. (A) SncRNA landscape revealed by traditional-seq; (B) SncRNA landscape revealed by PANDORA-seq. Pie charts depict the relative proportions of miRNAs, piRNAs, tsRNAs, rsRNAs, and other sncRNAs identified by each method. Bar graphs further illustrate the mean expression levels (RPM) of these sncRNA types across different size ranges. The error bars stand for the standard error of the mean. (traditional-seq n = 14 mice, PANDORA-seq n=16 mice)

### 3.3 Pronounced alteration of mitochondrial tsRNAs and rsRNAs in the prefrontal cortex of AD mice

To further understand the role of tsRNA and rsRNA in AD progression, we quantified their expression level in AD mice relative to WT counterparts, using both traditional-seq and PANDORA-seq data (Fig. 3). No significantly differentially expressed tsRNAs and rsRNAs were observed between WT and AD mice groups in the traditional-seq data (Fig. 3A), indicating that traditional-seq is not a sensitive method to detect the tsRNAs and rsRNAs profiles. In contrast, PANDORA-seq (Fig. 3B) detected numerous differentially expressed tsRNAs and rsRNAs, including 187 mitochondrial tsRNAs and 111 mitochondrial rsRNAs, which are all downregulated in the prefrontal cortex of the 5XFAD mouse brain. The genomic tsRNAs and rsRNAs displayed mixed expression patterns, with 43 decreased and 16 increased genomic tsRNAs, and 2,518 decreased and 84 increased genomic rsRNAs (Fig. 3B). Although the number of dysregulated genomic sncRNAs is greater than that of mitochondrial sncRNAs, Fisher’s exact test indicates that there is a significant enrichment of dysregulated mitochondrial tsRNAs (odds ratio = 18.176) and rsRNAs (odds ratio = 2.025) compared with the dysregulated genomic tsRNAs and rsRNAs, respectively. Consistent with the Fisher’s exact test results, further distribution analysis showed that the peaks for mitochondrial tsRNAs (Fig. 3C) and rsRNAs (Fig. 3D) are shifted further away from the center (log_2_FC = 0) compared to their genomic counterparts, indicating a greater degree of dysregulation in the AD cortex. Additionally, both mitochondrial sncRNA peaks were shifted to the left of the zero point, suggesting mitochondrial tsRNAs and rsRNAs are predominantly downregulated in AD (Fig. 3C and 3D).

**Fig. 3.**
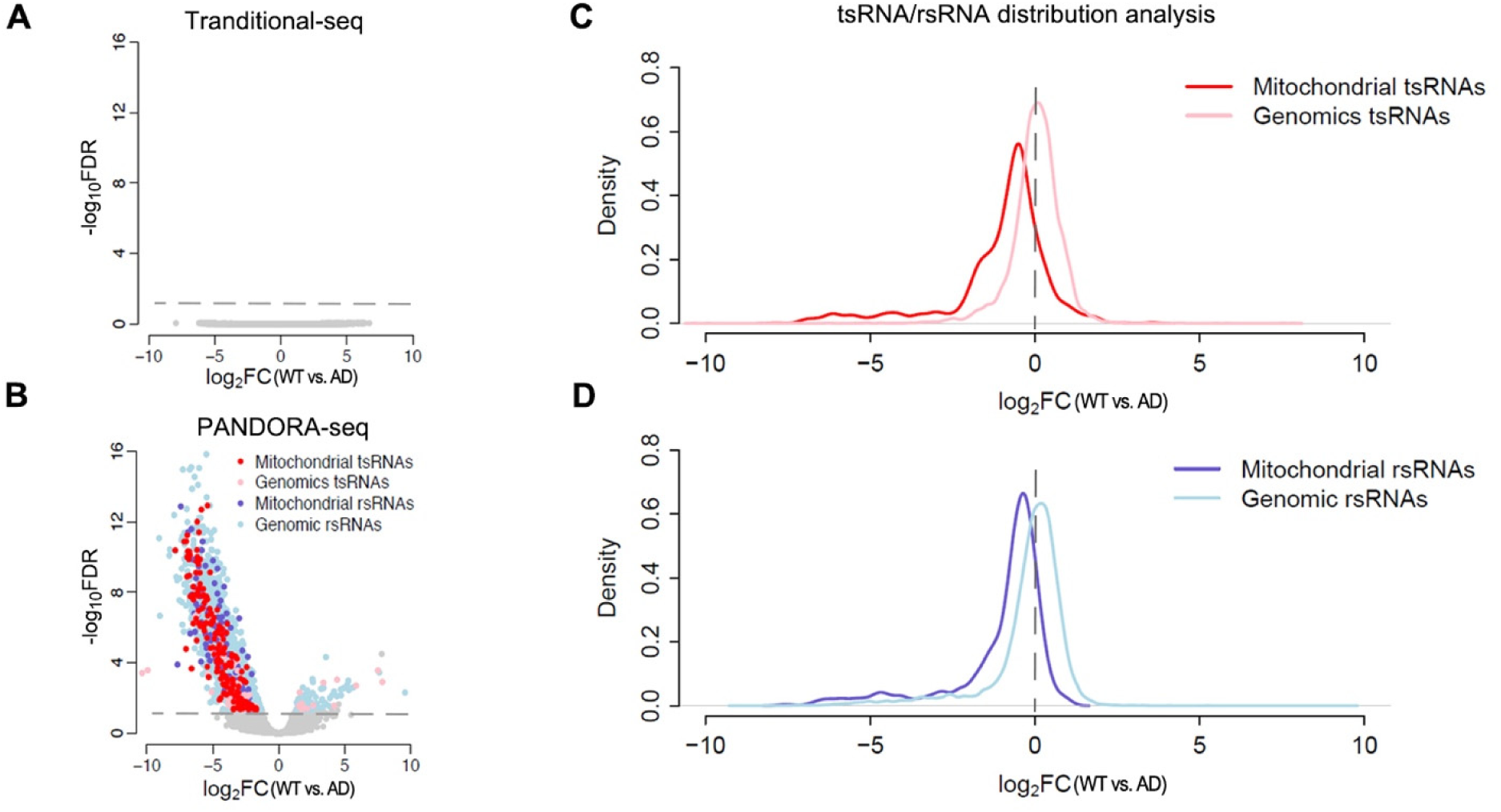
PANDORA-seq identified mitochondrial tsRNA/rsRNA as most sensitively dysregulated in AD mice. (A, B) Volcano plots of differentially expressed sncRNAs (DE-sncRNA) in the cortex of AD mice as compared with WT mice, revealed by traditional-seq (A) and PANDORA-seq (B). Colored dots represent the increased or decreased DE-sncRNAs with an *FDR*<0.05 and *FC*>2 as a cutoff threshold. (C, D) Probability density plots show the distribution of the log_2_ fold change (log_2_FC) of tsRNAs (C) and rsRNAs (D) in WT versus AD mice. (traditional-seq n = 7 mice in each group, PANDORA-seq n= 8 mice in each group)

### 3.4 Mitochondrial tsRNAs/rsRNAs are downregulated and negatively associated with lysosome pathway expression in AD cortex

TsRNAs and rsRNAs have been reported to play essential roles in various cellular activities, including, but not limited to, stem cell maintenance, cancer progression, epigenetic inheritance, and neurological diseases^27^. In this study, mitochondrial tsRNAs and rsRNAs exhibited a distinct downregulation pattern compared to other sncRNAs in the AD mice, promoting us to analyze the potential association between mitochondrial tsRNAs/rsRNAs expression and AD progression. The mapping profile in Figure 4 shows that three mitochondrial tsRNA families—mt-tsRNA-Gln, mt-tsRNA-His, and mt-tsRNA-Leu—as well as one rsRNA family derived from mitochondrial 16S rRNA (rsRNA-16S), are significantly reduced in the AD cortex. The RPM of mt-tsRNA-Gln was more than two-fold higher in the WT than that in the AD mice, with the majority of mt-tsRNA-Gln coming from the 30-74nt region of mitochondrial tRNA-Gln (Fig. 4A). Similarly, mt-tsRNA-His shows a two-fold decrease in RPM in AD samples, predominantly mapping to the 25-72 nt region of mt-tRNA-His (Fig. 4B). The RPM of mt-tsRNA-Leu is 1.5-fold lower in AD group, with mt-tsRNA-Leu mainly derived from the 35-78nt region of mt-tRNA-Leu (Fig. 4C). We also found that the rsRNAs in mitochondrial rsRNA-16S family, mapping to the 98-nt to 141-nt region, are reduced in the prefrontal cortex of AD mice (Fig. 4D). We also examined the expression profiles of genomic tsRNA-Gln, tsRNA-His, and tsRNA-Leu families, which are the genomic counterparts of the three mitochondrial tsRNA families. In contrast, the expression levels of these genomic tsRNAs showed no difference between WT and AD mice (Supplementary Fig. 2). These findings suggest a potential disruption in mitochondrial tRNA/rRNA processing or stability in AD that affects the mitochondrial tsRNA and rsRNA biogenesis.

**Fig. 4.**
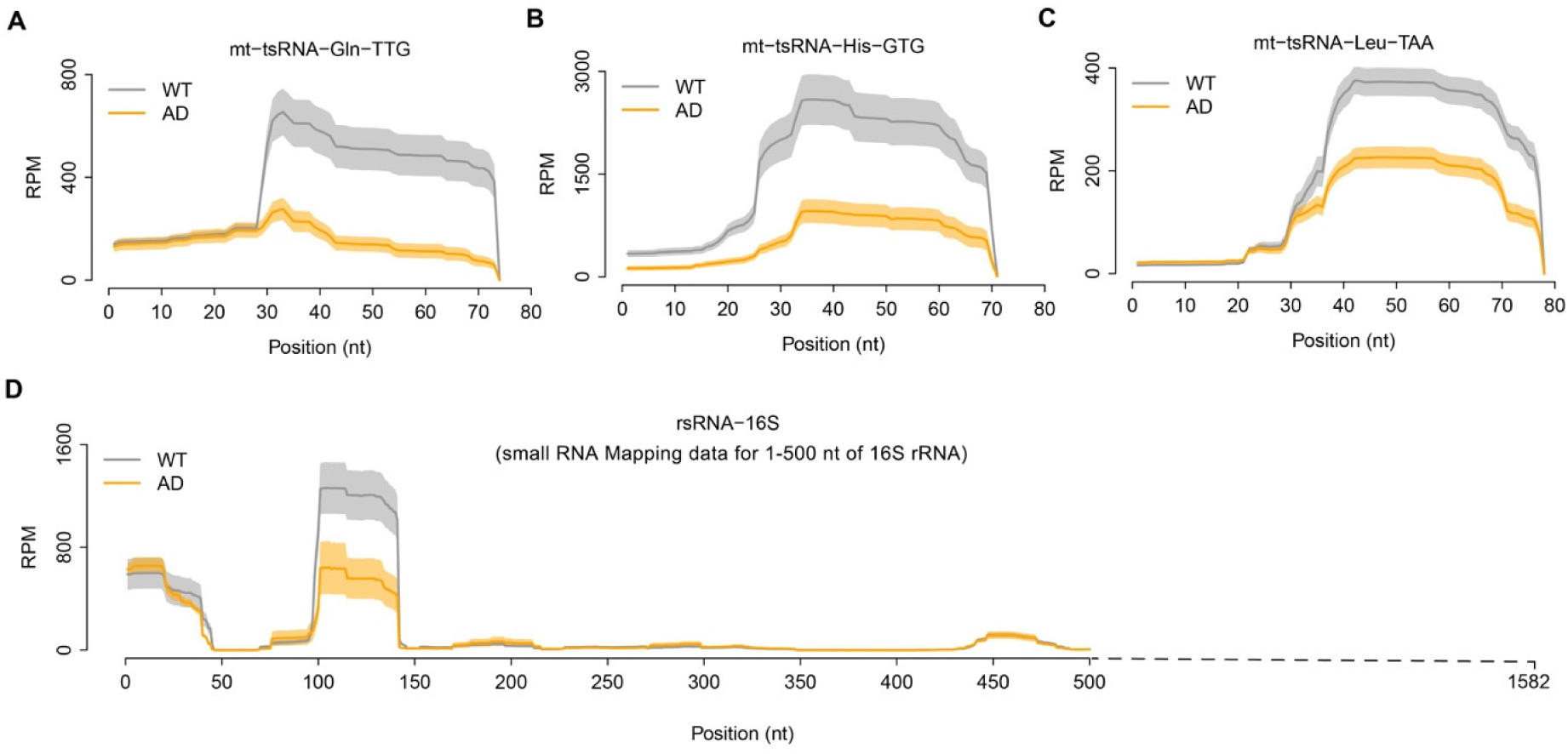
Mapping analyses of representative mitochondrial tsRNAs in WT and AD mice brain prefrontal cortex. (A) The expression profile of mt-tsRNA-Gln across the length in WT and AD samples. (B) The expression profile of mt-tsRNA-His across the length in WT and AD samples. (C) The expression profile of mt-tsRNA-Leu across the length in WT and AD samples. (D) The expression profile of mt-rsRNA-16S (101-140 nt) across the length in WT and AD samples. The solid curves indicate the mean of RPM, while the shading represents the standard error of the mean. (n = 8 mice in each group)

To explore how mitochondrial tsRNAs and rsRNAs potentially affect transcriptomic change and contribute to AD progression, we conducted transcriptome sequencing in WT and AD prefrontal cortex samples. A total of 191 differentially expressed genes (DEGs) were identified, with 179 of them upregulated in the AD group. Among these DEGs, certain genes associated with Alzheimer’s disease were identified, including Amyloid Beta Precursor Protein (APP), Presenilin 1 (PSEN1) and Cathepsin D (CTSD) *et al.*^28^ (Supplementary Fig. 3). Pathway enrichment analysis of these DEGs reveals that the lysosome pathway is the most significantly enriched (Fig. 5A). Lysosome-associated genes exhibit significantly higher expression in the AD cortex, with the heatmap revealing that all identified DEGs in lysosome pathway are exclusively upregulated. Notably, one outlier in the AD group showed comparable expression levels of the lysosome pathway to those in WT mice (Fig. 5B). These DEGs in the lysosome pathway are predominantly encoding various proteases responsible for protein degradation, exemplified by cathepsin (Cts) gene family and hexosaminidase (Hex) gene family. The co-expression analysis reveals significant negative correlations between lysosome gene expression and three prioritized mitochondrial tsRNA families and one rsRNA-16S family, with mt-tsRNA-Leu-TTA exhibiting the strongest negative association, as indicated by the correlation coefficient (ρ=-0.709) and the statistical significance (P = 0.003) (Fig. 5C). These findings underscore a potential regulatory link between mitochondrial tsRNAs and lysosome gene expression, which may contribute to the progression of Alzheimer’s disease.

**Fig. 5.**
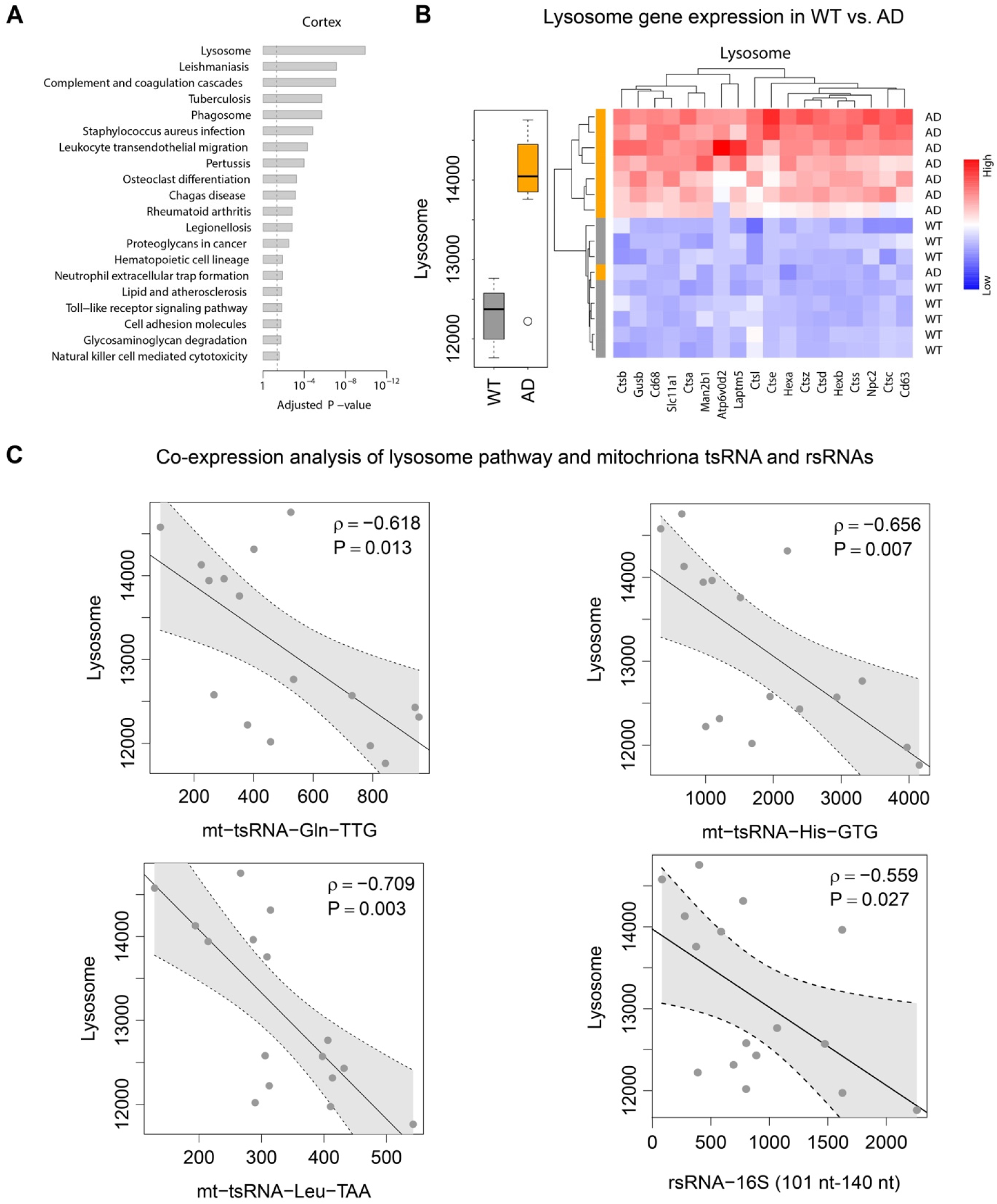
AD cortex shows increased lysosome activity, negatively correlated with mitochondrial tsRNAs. (A) KEGG pathway enrichment analysis of differentially expressed genes in WT versus AD cortex. The vertical dash line indicates the significance level of α=0.05. (B) The gene-set score of the lysosome pathway in WT and AD cortex (left); Heatmap representation of differentially expressed genes in the lysosome pathways (right). Red represents relatively upregulated expression, whereas blue represents reduced. (C) Relationship between the gene-set score of the lysosome pathway and the abundance of mt-tsRNA-Gln, mt-tsRNA-His, mt-tsRNA-Leu and mt-rsRNA-16S (101-140 nt). Each dot represents one sample (n=16). The grey band stands for the 95% confidence interval. P means statistical significance while ρ means the *Spearman*’s rank correlation coefficient. (n = 8 mice in each group)

## 4. Discussion

Emerging evidence has underscored the importance of small non-coding RNAs in various biological processes^29^, and the well-documented miRNAs and piRNAs, which regulate gene expression and protection of genomic integrity, have been shown to play critical roles in neurodegenerative disease^15, 30^. Recent studies also revealed the widespread presence and pivotal functions of novel classes of sncRNAs, such as tsRNAs and rsRNAs, in stress response, immunity, metabolism, and epigenetic inheritance of environmentally acquired traits^16, 27, 31, 32^. However, the functions of these novel sncRNAs in Alzheimer’s disease remain underexplored. In this study, we employed a widely used Alzheimer’s disease mouse model named 5XFAD transgenic mice and recently developed PANDORA-seq to explore the significance of tsRNAs/rsRNAs in AD progression.

The results of cognitive impairment test assays, including the Y-maze assay and NORT, showed 5XFAD mice have impairments of both short-term memory and recognition memory at the six-month stage. This is consistent with the saturation of Aβ plaques in the brain of 5XFAD mice by 6 to 9 months of age^4^. At two months old, when the Aβ plaques first appeared, the 5XFAD mice didn’t show any sign of cognitive impairment in the Y-maze assay and NORT. These outcomes confirm the efficacy of the 5XFAD mice model in mimicking clinical manifestations of Alzheimer’s disease.

To uncover the previously overlooked modified sncRNAs, we employed PANDORA-seq to comprehensively profile sncRNAs in the brain prefrontal cortex of WT and AD mice. PNDORA-seq uncovered a tsRNAs-and rsRNAs-enriched sncRNA landscape (Fig. 2B), which was strikingly different from the miRNA-enriched sncRNA landscape unveiled by the traditional sequencing method (Fig. 2A). Our results highlight the significant advantage of PADORA-seq in detecting modified sncRNA such tsRNAs and rsRNAs. This increased sensitivity for the detection of tsRNAs and rsRNAs also underscores the critical role that the modified sncRNAs may play in neural tissues.

The analysis of differentially expressed sncRNA showed that mitochondrial tsRNAs and rsRNAs were significantly downregulated in the cortex, while genomic tsRNAs and rsRNAs displayed mixed expression change between the WT and AD mice (Fig. 3A and Fig. 3B). The most significantly reduced mitochondrial tsRNAs are mt-tsRNA-Gln, mt-tsRNA-His, and mt-tsRNA-Leu in our study. We also identified some genomics tsRNA decreased in the AD mice, which aligns with our previous finding that genomic tsRNAs are downregulated in the post-mortem brain of human Alzheimer’s disease subjects^33^. Further transcriptome DEGs analysis and co-expression analysis showed that these reduced mitochondrial tsRNAs and rsRNAs are negatively associated with the expression of the lysosome pathway (Fig. 5). Mitochondria are known to play a crucial role in neuronal health, and their dysfunctions are connected to age-related neurodegenerative diseases, including Alzheimer’s disease and Parkison’s disease^34^. Lysosomes, essential for the degradation or recycling of biomacromolecules, play a significant role in cellular homeostasis, and their dysfunction is closely linked to the progression of Alzheimer’s disease^35^. The observed downregulation of mitochondrial tsRNAs/rsRNA and negative association with lysosome pathway in our analysis may suggest a scenario that enhanced lysosome activity in the AD cortex may lead to abnormal mitochondrial structure and function, with a decreased level of mitochondrial tsRNAs/rsRNA as either a molecular readout or the decrease of mitochondrial tsRNAs/rsRNA can further trigger mitochondrial and cellular events that promote the progression of Alzheimer’s disease. More importantly, the differential expression profile of mitochondrial tsRNAs has also been observed in other pathological conditions, including leukemia^36^, infection of pathogens^37^, oxidative stress^38^, and the high-diet exposed human sperm^39, 40^, underscoring the importance of mitochondrial tsRNAs in disease mechanisms.

The process underlying the biogenesis of mitochondrial tsRNA remains elusive despite many specific endoribonucleases involved in the production of genomic tsRNA having been identified in the cytosol^27, 41^. Unlike their cytosolic counterparts, no endoribonuclease has been definitively identified within mitochondria as responsible for cleaving mt-tRNA to generate specific mt-tsRNAs. RNase Z, known for trimming the 3’ trailer sequence of pre-tRNA in both mitochondria and nucleus^42^, has not been confirmed to cut mitochondria tRNA at other sites to produce mt-tsRNA. An intriguing hypothesis suggests that mt-tRNA may be exported to cytosol and processed by cytosolic endoribonucleases, supported by studies indicating the presence of mtRNA in the cytosol with RNA binding proteins (RBPs)^43, 44^. Moreover, specific modifications such as methylation and Queuosine (Q) on tRNAs are involved in the tsRNA production. For example, DNMT2- and NSUN2-dependent 5-methylcytosine (m^5^C) on tRNA increase its stability, making tRNA less vulnerable to Angiogenin (ANG)-mediated cleavage and consequently, tsRNA generation^9, 45, 46^. QTRT1-dependent queuosine (Q) modification can reduce the ANG-meditated tRNA cleavage and promote DNMT2-mediated m^5^C to further stabilize tRNA^47, 48^. Both NSUN2 and QTRT1 enzymes are found in the mitochondria^49, 50^, it is possible that different modification enzyme activities could influence mitochondria tsRNA production. Understanding the mt-tsRNA biogenesis can shed light on their potential role in modulating mitochondrial resilience or vulnerability under AD-associated stressors, offering insights into the complex interplay between mitochondrial function and neurodegenerative disease progression.

In summary, the current study utilized cutting-edge PANDORA-seq technique to reveal a distinctive mitochondrial tsRNA population implicated in the pathogenesis of Alzheimer’s disease. While the exact roles of these tsRNA in AD await discovery, their association with the disease underscores the potential for significant insights into their specific functions within this context. Future efforts to examine the biogenesis of mitochondria under pathological conditions would be invaluable and will open new avenues for dissecting the nature of dementia pathology and therapeutic strategies.

## Credit Authorship Contribution Statement

**Xudong Zhang:** Writing – review & editing, Writing – original draft, Methodology, Data curation, Investigation. **Junchao Shi**: Writing – review & editing, Investigation, Methodology. **Pratish Thakore:** Writing – review & editing, Data curation, Investigation, Methodology. **Albert L. Gonzales:** Writing – review & editing, Methodology. **Scott Earley:** Writing – review & editing, Conceptualization, Funding acquisition. **Qi Chen**: Writing – review & editing, Conceptualization, Funding acquisition. **Tong Zhou:** Writing – review & editing, Data curation, Formal Analysis, Conceptualization, Funding acquisition**. Yumei Feng Earley:** Writing – review & editing, Conceptualization, Data curation, Project administration, Funding acquisition.

## Data Availability

The sncRNA-Seq datasets have been deposited in the Gene Expression Omnibus (GSE277483). The sncRNA annotation pipeline SPORTS1.1 is available from GitHub (https://github.com/junchaoshi/sports1.1). Data supporting the plots within this article and other findings of this study are available from the corresponding author upon request.

## Acknowledgment

Small RNA library sequencing was conducted at the IGM Genomics Center, University of California, San Diego, La Jolla, CA.

---

**Supplementary Fig. 1.**
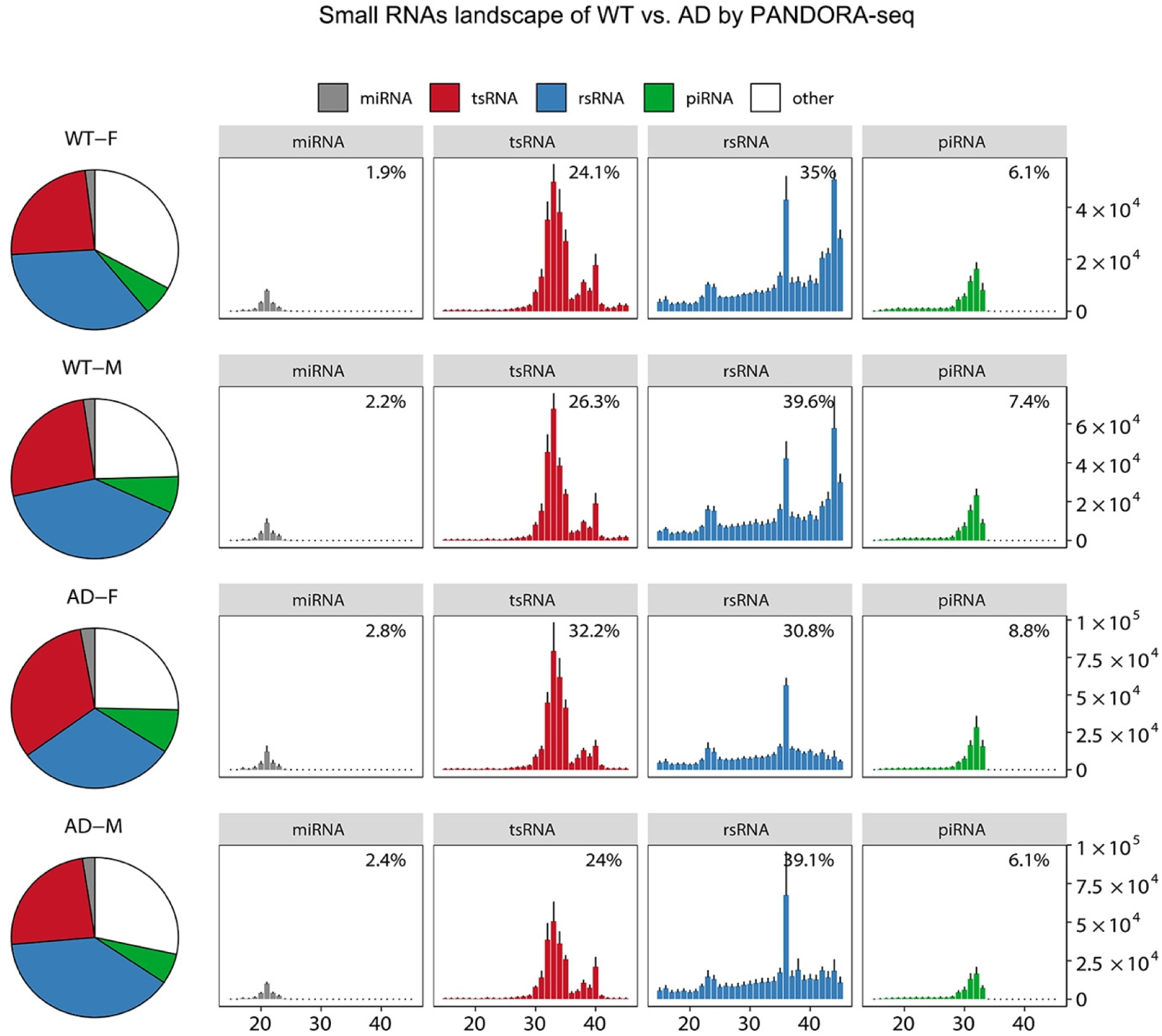
The small RNAs landscape of cortex from WT and AD mice detected by PANDORA-seq. Representative results of sncRNAs species detected by PANDORA-seq from WT and AD mice, including male and female. Pie charts depict the relative proportions of miRNAs, piRNAs, tsRNAs, rsRNAs, and other sncRNAs identified by each method. Bar graphs further illustrate the mean expression levels (RPM) of these sncRNA types across different size ranges. The error bars stand for the standard error of the mean. (n = 4 mice per sex in each group (WT and AD))

**Supplementary Fig. 2.**
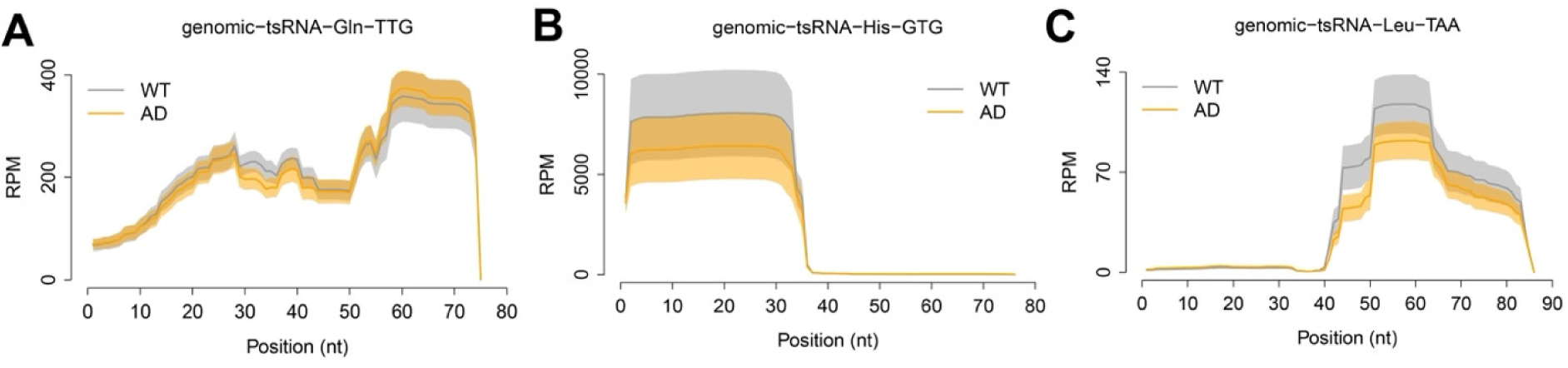
Mapping analyses of three genomics tsRNAs expression in WT and AD mice brain prefrontal cortex. (A) The expression profile of genomic tsRNA-Gln across the length in WT and AD samples. (B) The expression profile of genomics tsRNA-His across the length in WT and AD samples. (C) The expression profile of genomic tsRNA-Leu across the length in WT and AD samples. The solid curves indicate the mean of RPM, while the shading represents the standard error of the mean. (n = 8 mice in each group)

**Supplementary Fig. 3.**
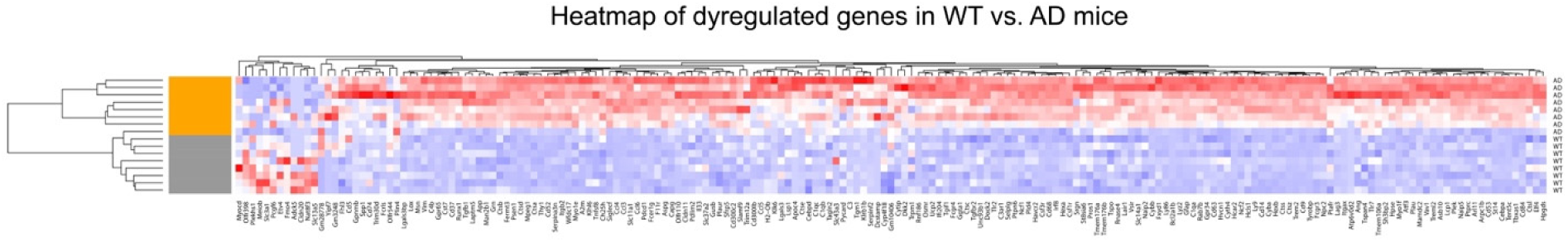
Transcriptomic change in the cortex between the WT and AD mice. Heatmap representation of differentially expressed genes in the cortex from WT and AD mice cortex. Red represents relatively upregulated expression, whereas blue represents reduced. (n = 8 mice in each group)

